# Mechanism-Based Discovery of New TRIDs Targeting Ribosome Pockets

**DOI:** 10.64898/2026.10.20.760543

**Authors:** Janeena Asuma, Hanfei Wang, Saleem Y. Bhat, Huile Zhu, Hong Li, Adam A. Jabak, David M. Chenoweth, Barry S. Cooperman

## Abstract

Approximately 10% of all human pathogenic mutations are nonsense mutations which give rise to premature termination codon (PTC) diseases, leading to an estimated 35 million people worldwide who harbor such mutations. To date, no approaches leading to medical treatments which increase PTC readthrough have received FDA approval. However, one well-tolerated translation readthrough inducing drug (TRID), ataluren, was conditionally approved for 11 years by the European Medicines Agency for treatment of nonsense mutation Duchenne Muscular Dystrophy. In recent work we have shown that ataluren increases readthrough indirectly, by exclusively inhibiting termination without directly affecting readthrough by suppressor aminoacyl-tRNAs (sup-aa-tRNAs). Based on the idea that this mode of action (MoA) might be the reason for ataluren’s low toxicity, we have discovered potential new TRIDs, which target pockets within the ribosome to which the termination factor eRF1 binds but sup-aa-tRNAs do not. These new TRIDs, dipeptide derivatives of eRF1 and some Sartans, mimic ataluren’s MoA. Three Sartans, which are angiotensin II receptor blockers and FDA-approved for treatment of hypertension, are significantly more potent than ataluren in our reconstituted *in vitro* readthrough assay. These compounds might, right now, be more effective than ataluren in treating some PTC diseases. We anticipate that further development of these lead compounds will lead to more general effectiveness, used both alone and in combination with other classes of readthrough inducers.

## Introduction

Over 7000 rare genetic diseases have been identified, collectively affecting an estimated 350 million people worldwide^**1**^. Approximately 10% of all human pathogenic mutations are nonsense mutations^**2**^ which give rise to premature termination codon (PTC) diseases, leading to an estimated 35 million people who harbor such mutations. This significant population would clearly benefit from treatments leading to increased PTC readthrough. Several therapeutic approaches are being pursued to overcome PTCs, as recently reviewed^**3–5**^. These include translation readthrough inducing drugs (TRIDs), notably including some which deplete cellular concentrations of eRF1^**6**,**7**^ or eRF3^**8**^, as well as nucleic acid-based approaches, including anticodon-edited suppressor tRNAs (ACE-tRNAs)^**9–11**^, mRNA and DNA editing and antisense oligonucleotides^**4**,**12**^. To date, none of these approaches has received FDA approval for treatment of any PTC disease.

One TRID, ataluren (Translarna), was conditionally approved for clinical use from 2014 - 2025 in treatment of nonsense mutation Duchenne Muscular Dystrophy (nmDMD) by the European Medicines Agency. The approval was withdrawn because the available evidence **“**failed to confirm that Translarna slowed disease progression or produced meaningful clinical improvement”. This despite ataluren’s very positive properties of **i**. showing consistent low toxicity in clinical trials for treatment of several different PTC diseases and **ii**. inhibiting termination at PTCs but not at NTCs (normal termination codons). These properties, and the abundant evidence that, in many cases, ataluren treatment benefits patients suffering from PTC diseases^**13–15**^, have spurred our efforts to determine ataluren’s mode of action (MoA) in promoting PTC readthrough, with the goal of developing new TRIDs which will be more generally effective than ataluren while preserving ataluren’s MoA. Here we report our recent progress toward achieving this goal.

In recent work, we have used a cell-free reconstituted system, denoted PURE-LITE (Fig. 1), to show that **i**. in contrast to toxic TRIDs like aminoglycosides, ataluren stimulates readthrough exclusively by competitively inhibiting release factor complex (RFC, eRF1.eRF3.GTP)-dependent catalysis of translation termination via binding to 3 or more sites in the protein synthesis machinery, without affecting readthrough of a PTC by Suppressor tRNA ternary complex (Sup-TC, Sup-aa-tRNA.eEF1A.GTP) in the absence of RFC^**16**,**17**^, and **ii**. ataluren’s ability to stimulate readthrough is negatively correlated with the strength of RFC interaction with the pretermination complex: i.e., ataluren is more stimulating when RFC interaction is weak and less stimulating when RFC interaction is strong^**18**^. These results have led us to hypothesize **i**. that ataluren’s MoA, in exclusively and reversibly inhibiting termination, may account for its low toxicity, and **ii**. that the variability of ataluren effectiveness in stimulating readthrough in clinical trials results from the dependence of the strength of RFC interaction with a PTC, a parameter which is highly sensitive to both the upstream and downstream mRNA contexts of the PTC.

**Fig. 1.**
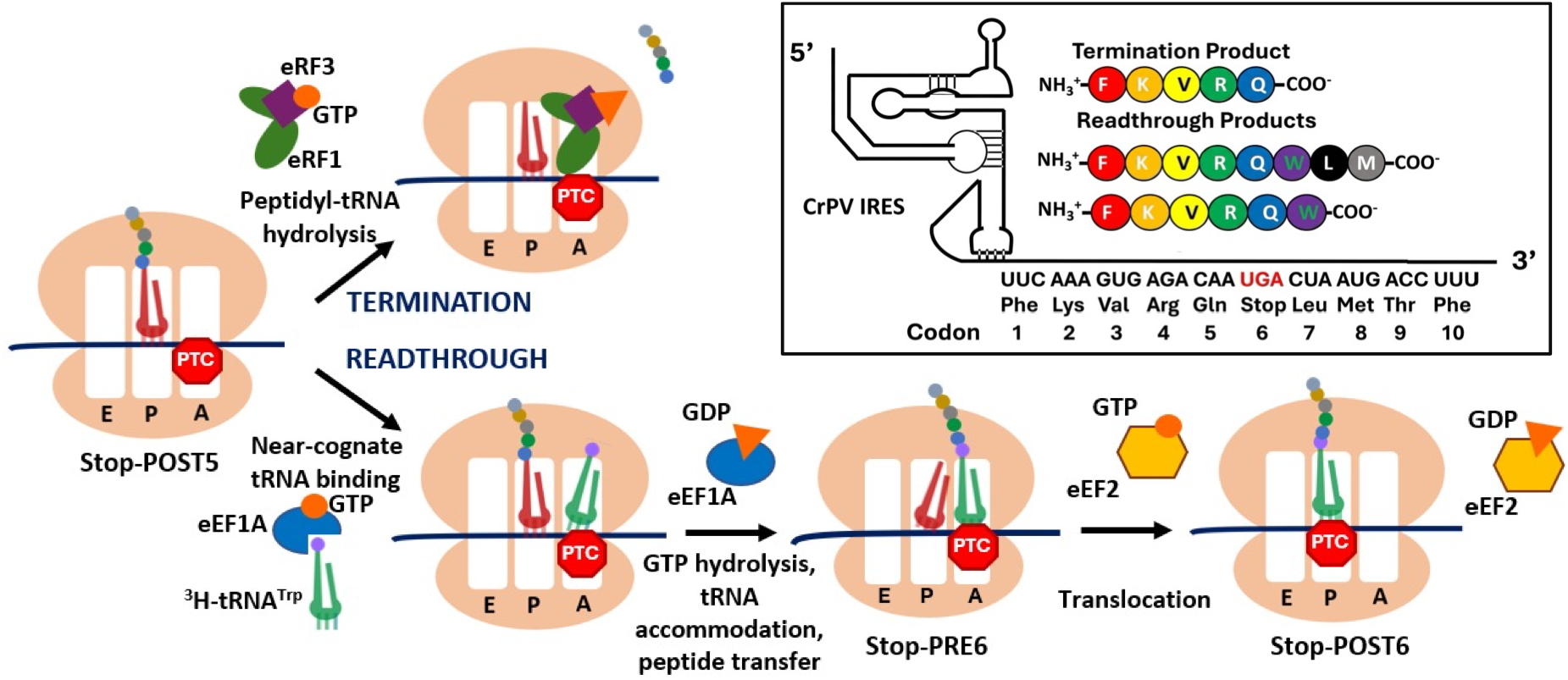
The PURE-LITE system for measuring termination and readthrough. RFC and Suppressor TC compete for reaction with Stop-POST5 pretermination complex. For readthrough of the reference mRNA sequence shown (inset), Trp-tRNA^Trp^ is used as suppressor aminoacyl-tRNA.

The structural basis for RFC competition with Sup-TC for binding to a posttranslocation complex (Fig. 1) is their molecular mimicry when bound to the ribosome^**24**^, with eRF3 mimicking eEF1A and eRF1 mimicking aa-tRNA. However, the latter mimicry is only partial: eRF1 has approximately twice the mass of aa-tRNA and, as a protein, can form interactions with spaces on the ribosome, which we denote as “eRF1 pockets”, not available to aa-tRNA. In this manuscript we describe two distinct classes of new TRIDs, oligopeptides, and some repurposed Sartans, a group of angiotensin II receptor blockers which are FDA-approved for treatment of hypertension. Both share with ataluren the ability to inhibit RFC catalysis of termination while not disturbing Sup-TC readthrough of a PTC.

In what follows we combine **i**. detailed comparison of published structures of A-site bound eRF1 and aa-tRNA to identify two distinct eRF1 pockets; **ii**. functional characterization of eRF1 variants at positions interacting directly with ribosome residues within eRF1 pockets; **iii**. TRID docking within eRF1 pockets and **iv**. determination of TRID activities in inhibiting termination and stimulating readthrough.

## Results

### Identification of eRF1 pockets

We used two criteria to identify two eRF1 pockets, denoted 40S-1, and 60S-1, which contain residues closely interacting with eRF1 residues (distances of closest approach <3.9 Å), but not with A-site bound aa-tRNA (distances of closest approach >5.9 Å) in the 40S and 60S subunits of mammalian ribosomes, respectively. In applying these criteria, we examined known cryo-EM structural models of the ribosome during termination, when it is bound to eRF1 during elongation (PDBs: 5LZU, 7NWH), and when it is bound to A-site aa-tRNA (PDB: 6Y57, 3J7R). Both eRF1 and aa-tRNA bind in two forms to the ribosome, a compact form when each initially bind as part of a complex, RFC (eRF1.eRF3.GTP) and TC (aa-tRNA.eEF1A.GTP), respectively, and an extended form, reaching into the peptidyl transferase center, after GTP is hydrolyzed and the chaperone protein nucleotide complex, eRF3.GDP and eEF1A.GDP, respectively, is released from the ribosome. We designate these forms as eRF1_C_ (compact, Fig. 2A), eRF1_E_ (extended) (Fig. 2B) and aa-tRNA_C_ (compact), aa-tRNA_E_ (extended), respectively. Distances of closest approach to ribosome residues in each of the two eRF1 pockets from eRF1_C,_ eRF1_E_, aa-tRNA_C_, and aa-tRNA_E_. are listed in Table 1. Some of the closest residues in pockets 60S-1 and 40S-1 are shown in red in Figs. 2C and 2D, respectively. Also shown in red in Fig. 2D is the A1240 residue, neighboring G1236 and C1237, which we earlier identified by photoaffinity labeling as part of a functionally important ataluren binding site^**17**^.

**Table 1.** Ribosome contact residues with specific eRFl sites and closest approach to A-site bound tRNAs.

| Table 1. Ribosome contact residues with specific eRF1 sites and closest approach to A-site bound tRNAs |  |  |  |  |  |
| --- | --- | --- | --- | --- | --- |
| eRF1 conformer, pocket, pdb | eRF1 residue | Ribosome residue | Closest Contact (Å) | Nearest tRNA<sub>E</sub> (Å)<br>pdbs:3J7R; 6Y57 | Nearest tRNA<sub>C</sub>(Å)<br>pdb:5LZS |
| eRF1<sub>E</sub>, 40S-1, 5LZU | T76 | G1236 (18S) | 3.657 | 13.384 | 14.097 |
|  | S77 | C1237 (18S) | 3.880 | 13.586 | 14.665 |
|  | Q80 | G1236 (18S) | 3.036 | 13.384 | 14.097 |
| eRF1<sub>E</sub>, 60S-1, 5LZU | F385 | G1981 (28S) | 3.150 | 39.535 | 5.870 |
|  | R414 | G1981 (28S) | 2.999 | 39.535 | 5.870 |
|  | Y415 | A2009 (28S) | 2.973 | 49.764 | 7.103 |
| eRF1<sub>C</sub>, 60S-1, 7NWH | K310 | G1981 (28S)<br>A2009 (28S) | 1.936<br>2.324 | 39.535<br>46.374 | 5.870<br>7.103 |
|  | E313 | G1981 (28S) | 2.362 | 39.535 | 5.870 |
|  | M314 | A2009 (28S) | 2.149 | 46.374 | 7.103 |
|  | G386 | U4419 (28S)<br>U4420 (28S) | 2.136<br>2.845 | 34.538<br>36.089 | 11.351<br>16.111 |
|  | A387 | U4419 (28S) | 3.034 | 34.538 | 11.351 |
|  | T388 | U4419 (28S) | 2.770 | 34.538 | 11.351 |
|  | R414 | U4419 (28S)<br>U4420 (28S) | 2.077<br>1.902 | 34.538<br>36.089 | 11.351<br>16.111 |
|  | Y415 | U2008 (28S)<br>A2009 (28S) | 1.955<br>1.435 | 49.764<br>46.374 | 14.000<br>7.103 |
|  | R416 | U2008 (28S) | 2.199 | 49.764 | 14.000 |
|  | V417 | U2008 (28S)<br>A2009 (28S) | 2.266<br>2.303 | 49.764<br>46.374 | 14.000<br>7.103 |
|  | D418 | U2008 (28S)<br>A2012 (28S) | 3.069<br>2.976 | 49.764<br>56.551 | 14.000<br>18.746 |
|  | M422 | U2008 (28S)<br>A2009 (28S)<br>C2011 (28S) | 2.742<br>2.185<br>1.844 | 49.764<br>46.374<br>53.072 | 14.000<br>7.103<br>14.450 |
|  | E423 | A2010 (28S) | 2.008 | 48.448 | 9.488 |
|  | Y424 | A2009 (28S)<br>A2010 (28S)<br>A1984 (28S) | 2.206<br>2.330<br>2.354 | 46.374<br>48.448<br>49.498 | 7.103<br>9.488<br>16.446 |

**Figure 2.**
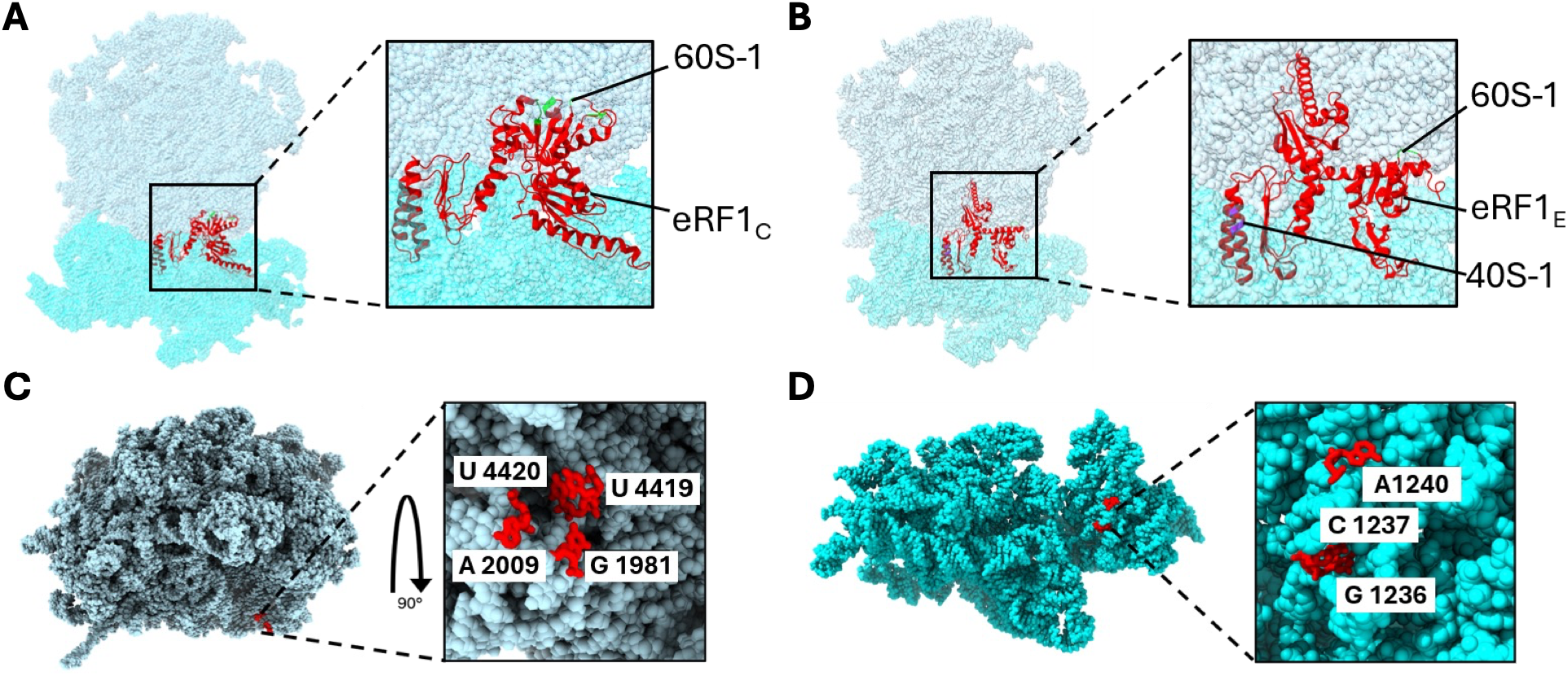
**A-B**. Structural models of eRF1 bound to the mammalian ribosome with locations of eRF1 pockets indicated. **A**, eRF1_c_ bound, pdb 7NWH. **B**. eRF1_E_ bound, pdb 5LZU. The 60S subunit is in blue, the 40S subunit is in cyan, and eRF1 is in red. Note that the 40S-1 pocket only becomes evident when eRF1 is elongated. **C, D**. Ribosome nucleotides (in red) flanking 60S-1 and 40S-1, PDBs 3J7R and 6Y57, respectively. A1240 is adjacent and was previously identified by photoaffinity labeling as being proximal to an ataluren binding site^17^.

### Functional characterization of eRF1 variants

Measuring termination activity at a PTC as a function of RFC concentration allows calculation of the parameters K_m_^RFC^ and k_cat_^**18**^. We used site-specific mutation to prepare and purify 31 variants of human eRF1 and determined the effects of each mutation on these parameters (Tables S1,S2). Variants in six eRF1 residues, three directly interacting with each of the eRF1 pockets, lead to 2 – 10 fold increases in K_m_^RFC^, with smaller changes in k_cat,_ values (Table 2).

**Table 2.** Mutation effects on termination kinetics.

| **Table 2.** Mutation effects on termination kinetics |  |  |  |  |
| --- | --- | --- | --- | --- |
| | eRF1 variant | WT-eRF1 contact with rRNA | $K_m^{RFC}$, nM | $k_{cat}$, min<sup>-1</sup> |
| eRF1 pocket | Wild Type | G1236<br>C1237<br>G1981<br>U2008<br>U4419<br>U4420 | $10 \pm 2$ | $0.30 \pm 0.02$ |
| 60S-1 | T388A | U4419 | $23 \pm 4$ | $0.18 \pm 0.01$ |
| | R414A | G1981,<br>U4419,<br>U4420 | $85 \pm 24$ | $0.22 \pm 0.04$ |
| | R416A | U2008 | $41 \pm 9$ | $0.35 \pm 0.02$ |
| 40S-1 | T76A | G1236 | $104 \pm 11$ | $0.27 \pm 0.02$ |
| | S77E | C1237 | $51 \pm 8$ | $0.19 \pm 0.02$ |
| | Q80A | G1236 | $77 \pm 10$ | $0.22 \pm 0.02$ |

These results suggest that eRF1 interaction with each of the eRF1 pockets, as measured by K_m_^RFC^, contributes to the overall strength of RFC functional binding. In contrast, the lesser effects on k_cat_ values indicate that these mutations have little effect on the structural integrity of eRF1, given the generally extreme sensitivity of k_cat_ values to even minor perturbations of enzyme structure. In support of including S77 as important for eRF1 binding to 40S-1, the S77E variant has recently been shown by others^**20**^ to increase readthrough of all three stop codons in 12 pathogenic PTC reporters in multiple disease contexts.

### Activities of oligopeptide TRIDS as termination inhibitors and readthrough stimulators

The presence of consecutive amino acids in eRF1 which interact with eRF1 pockets (Tables 1 and 2) suggested to us that oligopeptides mimicking these residues, 2-8 amino acids in length, might interfere with RFC function on the ribosome with effects on termination and readthrough similar to those of ataluren. We have so far tested some 30 oligopeptide and oligopeptide derivatives modified at one or both N- and C-termini (Fig. S3). The most successful to date are the termini-protected derivatives of the dipeptides R^414^Y^415^, N-AcRY-OMe, denoted RY-1, N-Ac-(4-CF_3_)-F-OMe, denoted RY-2, and T^76^S^77^, N-AcTS-OMe, denoted TS-1. As shown in Fig. 3 and summarized in Table 3, RY-1, RY-2, and TS-1 act similarly to ataluren, with comparable activities in inhibiting termination, and stimulating readthrough activity in the presence of RFC, but not in its absence. The derivatized dipeptide N-Ac-YR-OMe had negligible activity in both assays, an indication of the structural specificity required for such activities.

**Table 3.** TRID Termination and Readthrough EC_50_S.

| **Table 3. TRID Termination and Readthrough** |  |  |
| --- | --- | --- |
| EC<sub>50</sub>s |  |  |
| TRID | Termination<br>EC<sub>50</sub> (mM) | Readthrough<br>EC<sub>50</sub> (mM) |
| Ataluren | 0.32 ± 0.04 | 0.36 ± 0.03 |
| TS-1 | 0.72 ± 0.05 | 0.63 ± 0.05 |
| RY-1 | 0.55 ± 0.05 | 0.52 ± 0.08 |
| RY-2 | 0.40 ± 0.03 | 0.40 ± 0.06 |
| Telmisartan | 0.06 ± 0.01 | 0.038 ± 0.006 |
| Candesartan | 0.09 ± 0.01 | 0.074 ± 0.015 |
| Candesartan<br>Cilexetil | 0.004 ± 0.001 | 0.006 ± 0.002 |

**Fig. 3.**
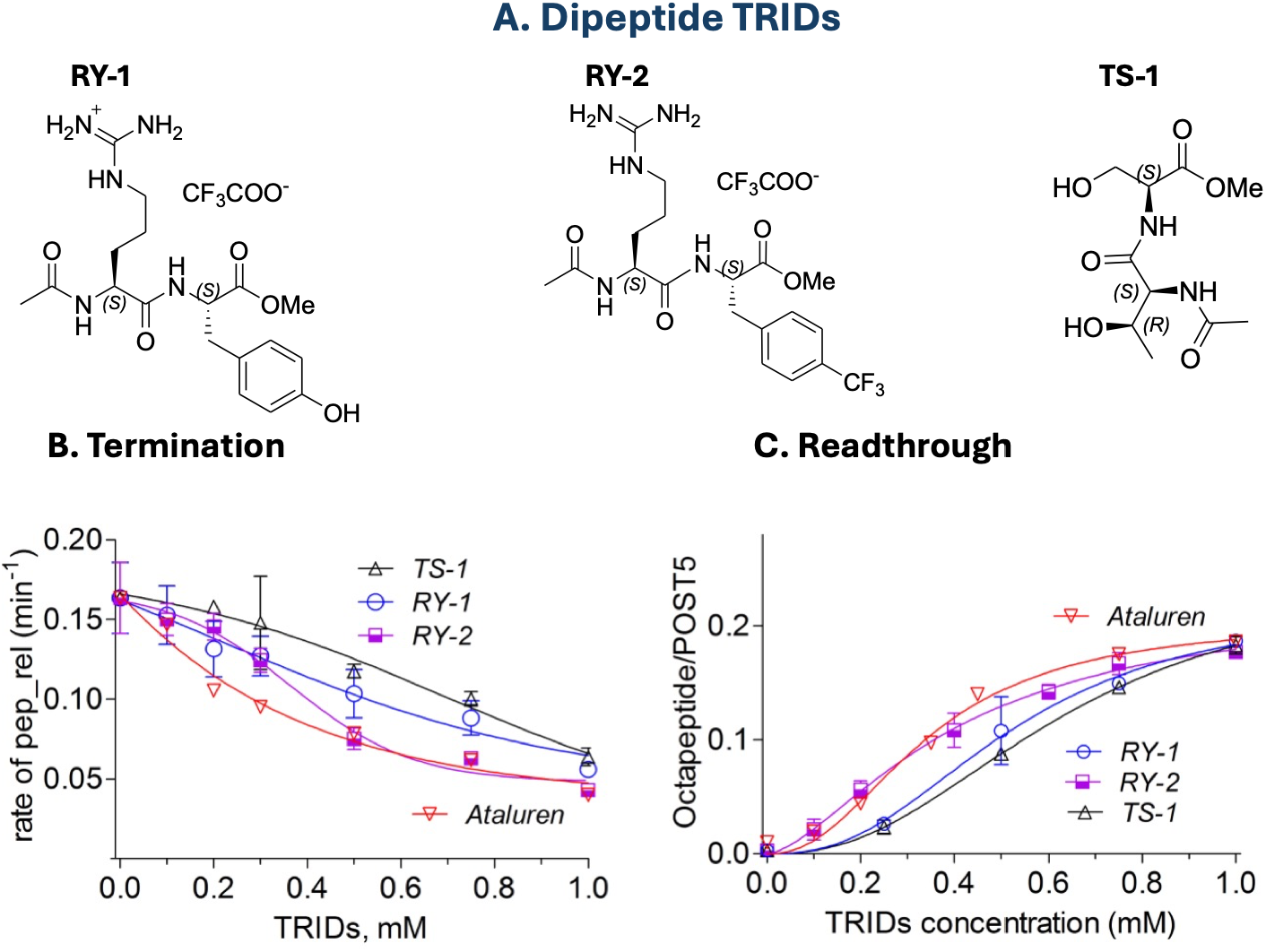
Activities of dipeptide TRIDs in inhibiting termination and inducing readthrough in PURE-LITE assays.

### Docking of RY-1 and TS-1

Due to its relatively high compatibility with nucleic acids, we used rDock to identify possible binding motifs of RY-1 and TS-1 to the 60S-1 and 40S-1 eRF1 pockets, respectively, as described in detail in Materials and Methods. For docking to 60S-1 we targeted spheres of 15 Å radius from N9 of A2009 (60S-1 pocket, pdb 3J7R) and N1 of 1237C (40S-1 pocket, pdb 6Y57) within the human ribosome structural models. In both cases, the highest scoring poses always led to docking of RY-1 within 60S-1 and TS-1 within 40S-1 with only minor variation, resulting in close contacts (≤ 3.2 Å) with a limited set of rRNA nucleotides (Fig. 4, Tables S3, S4), including those in contact with eRF1 residues shown to be important for eRF1 termination activity (Table 2).

**Figure 4.**
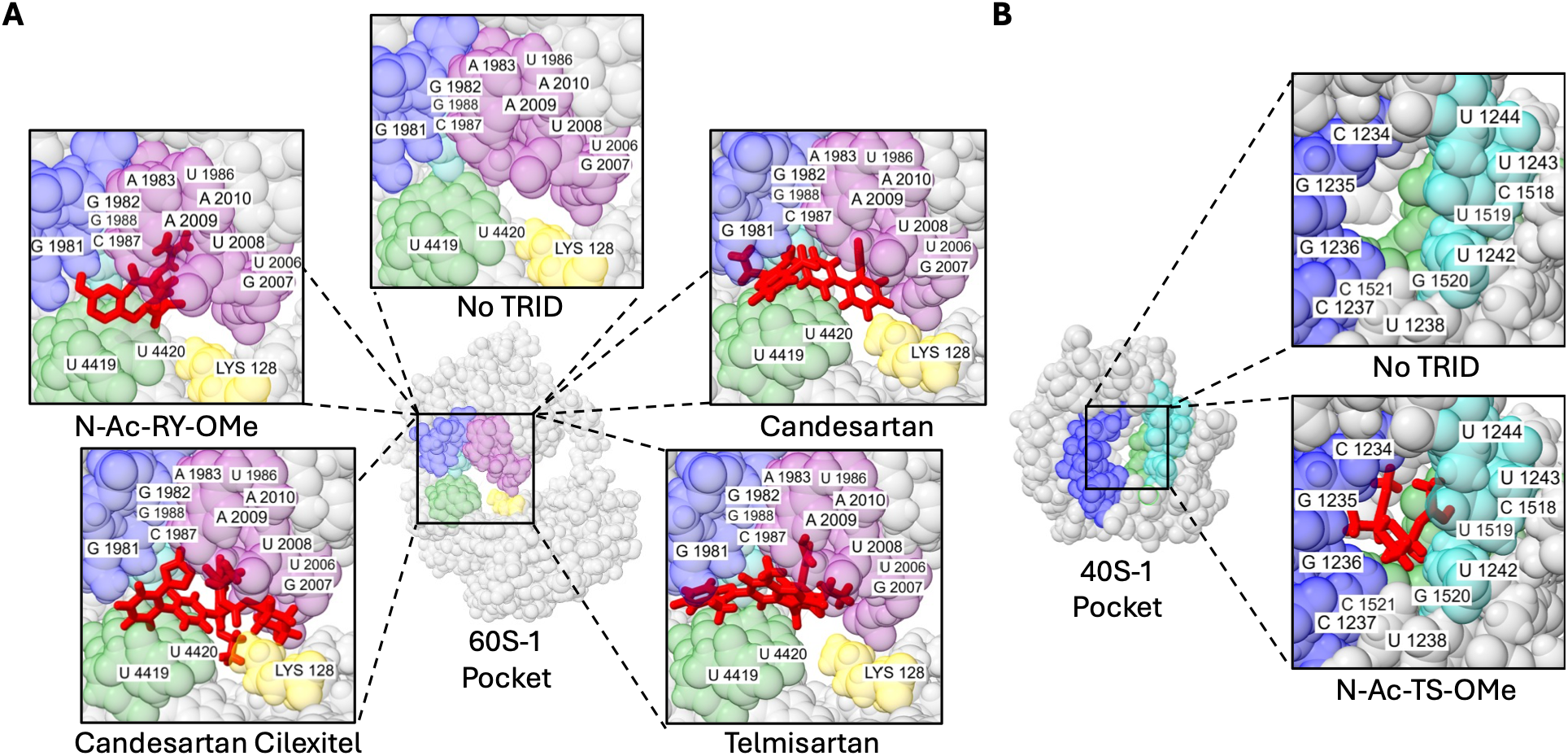
Best docking poses for N-Ac-RY-OMe, Telmisartan, Candesartan and Candesartan Cilexitel to the 60S-1 pocket (**A**) and for N-Ac-TS-OMe to the 40S-1 pocket (**B**). In **A** and **B**, different colors, four and two, respectively, are used to highlight different contact regions in each of the two pockets.

### High throughput virtual screening targeting the 60S-1 eRF1 pocket, leading to the identification of potential TRIDs having activity similar to ataluren but with significantly higher activity

The large number of eRF1:ribosome contacts contained within eRF1 pocket 60S-1 (Table 1) led us to prioritize targeting the same sphere of 15 Å radius from N9 of A2009 as above in performing a high throughput virtual docking screen of the CSD Drug Subset^**21**^, a collection containing published small-molecule crystal structures of ~ 17,000 clinically approved drug molecules. We initially identified over 20 high scoring candidate molecules of which we selected four, sulfamethoxazole, droperidol, diosmin, and telmisartan, as especially promising. We next compared the activities of these four compounds as termination inhibitors with that of ataluren. Measured at 300 µM, telmisartan was a much stronger inhibitor than ataluren, whereas the other three showed little or no activity. This result prompted us to determine the termination activities of other commercially available Sartans, a class of structurally related angiotensin II receptor blockers. These included candesartan, candesartan cilexetil, losartan, valsartan, olmesartan, aprosartan, and irbesartan. The preliminary results (Fig. S2) showed high inhibition activities for candesartan and candesartan cilexetil, an activity for losartan comparable to that of ataluren, and much lower activities for the others.

### Activities of Sartan TRIDS as termination inhibitors and readthrough stimulators

We followed up these promising results with more detailed examination of the termination and readthrough activities of the three most active Sartans. Their higher activities in the PURE-LITE assays of termination inhibition and readthrough stimulation assays compared with those of ataluren are displayed in Fig. 5 and Table 3. The values presented in Table 3 show that their EC_50_s for termination and readthrough stimulation range from 4 − 80 fold lower to 5 − 60 fold lower, respectively, with relative potency decreasing in the order candesartan cilexetil >> telmisartan >candesartan.

**Fig. 5.**
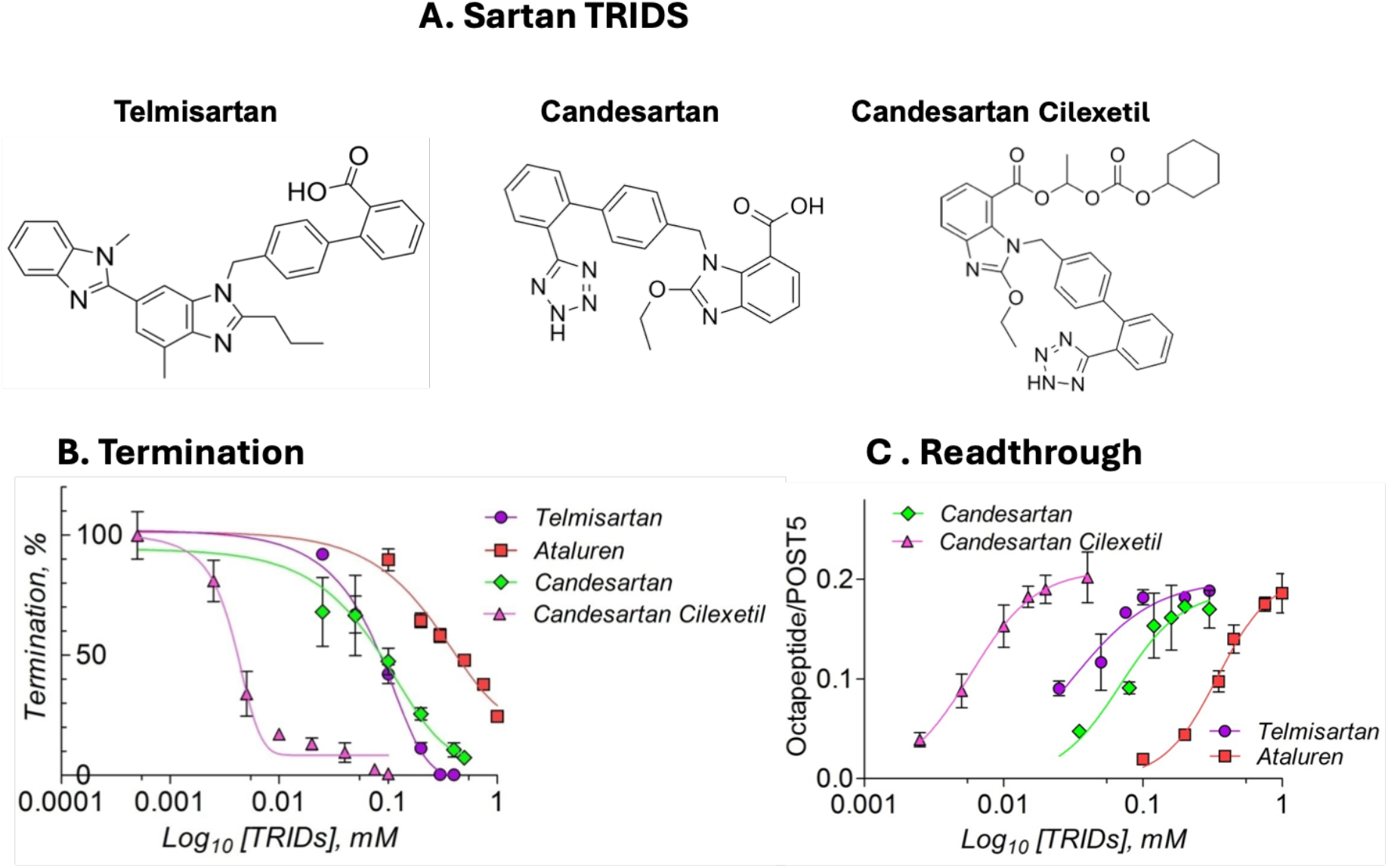
Activities of Sartan TRIDs in inhibiting termination and inducing readthrough in PURE-LITE assays.

We evaluated the ability of these Sartans to promote readthrough in a more biologically relevant assay by determining their activities in stimulating readthrough in rabbit reticulocyte extract of PTCs falling within CFTR (G542X, R1162X, W1282X), and Fibrillin1 (R2694X), which give rise to cystic fibrosis and Marfan syndrome, respectively. For this purpose we used a dual reporter assay (Fig.6A), in which genes encoding two different luciferases are placed upstream and downstream of an inserted 153 nt mutant DNA encoding 75 nucleotides both upstream and downstream of the in-frame UGA codon, providing a full sequence context for the PTCs listed above. The results (Fig. 6B-D, Table 4) demonstrate that each of the Sartans induce significant readthroughs, often going through a maximum as a function of Sartan concentration. Overall, induced readthrough, evaluated as observed readthrough in the presence of a Sartan minus readthrough in the absence of Sartan, was most pronounced for R1162X and R2694X, lesser for G542X and almost negligible for W1282X. We think it likely that the reduced activity of the prodrug candesartan cilexetil in the reticulate extract assay compared to what we observed in the PURE-LITE assays (Table 3), similar to that of candesartan, is due to esterase activity, a possibility that requires confirmation. From our PURE-LITE results we expected telmisartan and candesartan to induce similar readthroughs, irrespective of which PTC they targeted. This was true for FBN1-R2694X, but not for either CFTR-R542X or CFTR-1162X, for which telmisartan and candesartan have clearly different readthrough efficiencies. This observation merits further clarification. One possibility, which could account for these differences, is that each of these Sartans may be able to bind to the decoding center of an empty A-site, thus interfering with readthrough, in a distinct mRNA-sequence dependent manner.

**Table 4.** Induced readthrough measured in rabbit reticulocyte extract with basal readthrough subtracted^a^.

|  |  |  |  |  |
| --- | --- | --- | --- | --- |
| PTC | Concentration at maximum readthrough |  |  | Neighboring sequence context |
|  | Telmisartan | Candesartan | Candesartan Cilexetil |  |
| CFTR-G542X | 5.6 ± 1.0 %, 10 μM | 1.3 ± 0.1%, 50 μM | 2.1 ± 0.4%, 25 μM | CUU **UGA** GAA GGU |
| CFTR-R1162X | 6.1 ± 0.4%, 100 μM | 7.6 ± 1.2%, 10 μM | 6.8 ± 0.5%, 10 μM | AGC **UGA** GUC UUU |
| FBN1-R2694X | 8.5 ± 0.1% , 50 μM | 8.6 ± 1.9%, 10 μM | 4.2 ± 0.1%, 50 μM | GGC **UGA** GGAAAC |

<sup>a</sup>Basal readthrough is readthrough in absence of added Sartan; values are ± SD.

**Fig. 6.**
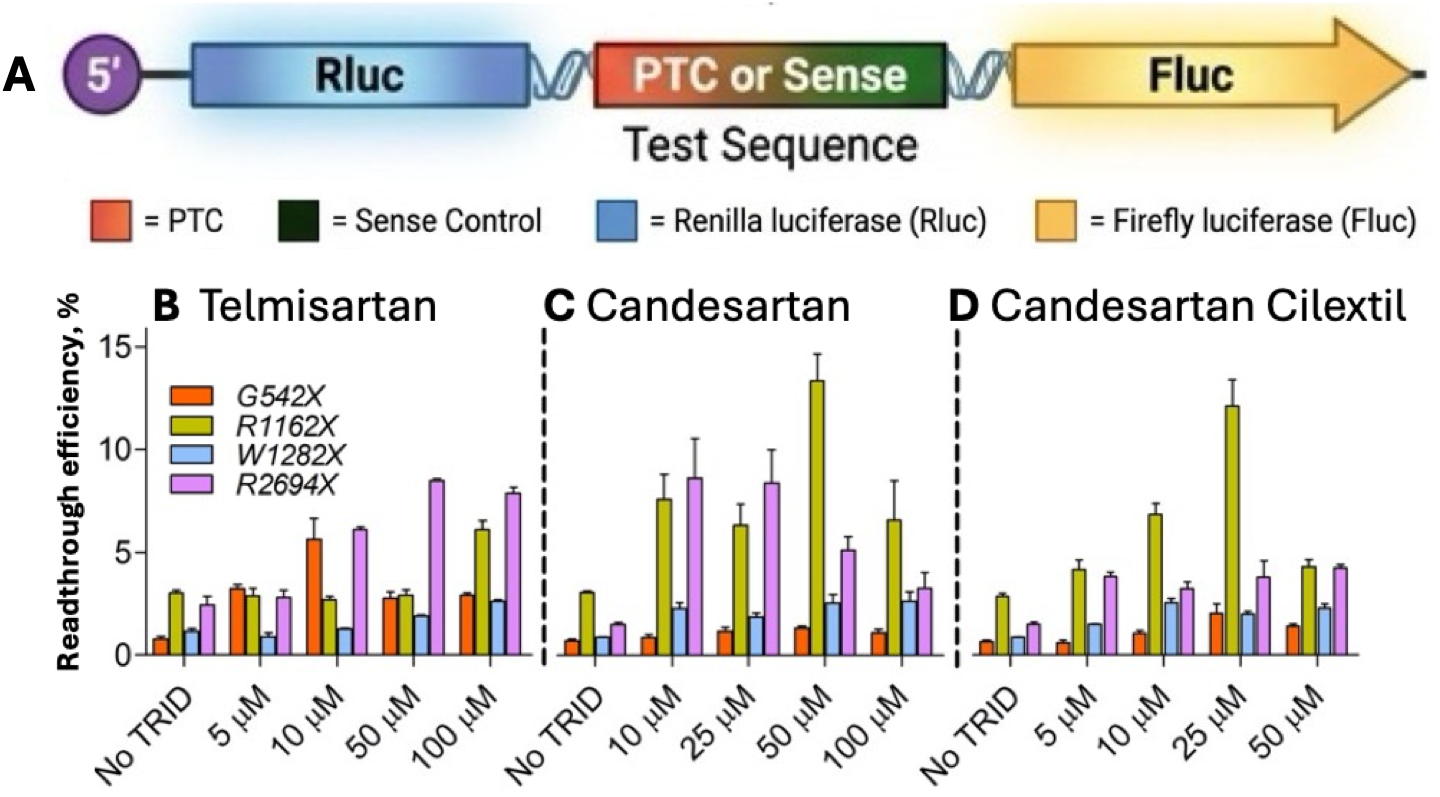
Activities of Telmisatan, Candesartan, and Candesartan Cilexetil in stimulating readthrough of CFTR PTCs G542X, R1162X, and W1282X and FBN1 PTC R2694X at the indicated concentrations (5-100 pM). No TRID represents the untreated solvent control. Bars show mean readthrough efficiency (%) ± SEM (n = 4-6). Dashed vertical lines separate the three compounds.

## Discussion

Here we have shown that identifying and targeting what we have defined as eRF1 pockets in the mammalian ribosome has led us to discover two distinct classes of TRIDs, dipeptides and Sartans, which precisely mimic ataluren in specifically inhibiting RFC termination activity while not directly affecting suppressor TC activity. Both classes have potential for increasing readthrough in patients harboring penetrant PTC mutations. This potential is currently more evident for the Sartans, which in addition to being considerably more potent than ataluren in stimulating readthrough (Table 3, Fig. 5) have, as repurposed FDA-approved drugs for treating hypertension, low toxicity. ***As a result, we believe it likely that they may already be capable of improving the treatment of a greater proportion of PTC patients than has been true for ataluren***. We stress this point, because of the significant number of patients afflicted by PTC diseases who have benefited from ataluren treatment^r**13–15,22,23**^ (see also Table S6).

We are clear in our understanding that our present results represent only an important first step towards the goal of discovering even more potent TRIDs capable of more successful general treatment of PTCs, used either alone or in combination with nonsense suppressors acting by other mechanisms, such as anticodon-edited tRNAs^**19**^. Highlighting our present limited knowledge is our lack of definitive structural evidence for the hypothesis that the 60S-1 and 40S-1 pockets actually function as TRID binding sites. With a collaborator, we are currently seeking such evidence. Nevertheless, the results demonstrating the sensitivity of the eRF1 binding activity to mutation of specific eRF1 residues flanking the pockets (Table 2) strongly supports this hypothesis. Also in support of the 40S-1 pocket as the dipeptide binding site is our previous identification by photoaffinity labeling of A1240, which neighbors 18S rRNA residues 1236-1238 and 1242 flanking the 40S-1 pocket (Table 2, Fig. 2D).

In work currently underway we, are conducting additional high throughput virtual screens to identify new TRIDs specific for binding to the 40S-1 pocket. We also are building on the result from our PURE-LITE assays that the ester, candesartan cilexetil, has ~ 10-fold greater activity in inducing readthrough than candesartan itself (Fig. 5, Table 3), to determine whether optimization of esterase-resistant derivatives of the carboxylic acids of both candesartan and telmisartan will yield the more powerful TRIDs which we seek. Finally, we are exploring the possibility of identifying additional eRF1 pockets that will bind to new TRIDs.

## Materials and Methods

### Identification of ribosomal sites/pockets

The identification of sites was performed using ChimeraX and cryo-EM structural models of the rabbit reticulocyte ribosome with eRF1 bound (PDB: 5LZU) in extended conformation from Shao *et al*.^**24**^ as well as with the eRF1 bound in compact conformation and eRF3 bound (PDB: 7NWH).^**25**^ All of the contacts within 4 Å between eRF1 and the 40S as well as the 60S ribosomal subunits were identified using the contacts function of ChimeraX. The rRNA nucleotides involved were flagged for measurement of distance from tRNA during elongation, using the distance function of ChimeraX. The distances between the tRNA in an A/P hybrid state (PDB: 6Y57 for 40S-1 and 3J7R for 60S-1 respectively) and the identified rRNA nucleotides at the nearest C1’ atoms of the respective species were measured using the distance function, and all of the rRNA nucleotides which interacted with eRF1 but did not interact with tRNA (defined by a distance between the nearest C1’ atoms no less than 5 Å) were included for consideration as potential sites for targeting with peptides and small molecules by virtual screen.

### Docking of ligands within eRF1 pockets

The receptors used for docking were fragments of ribosomal PDBs 6Y57 and 3J7R centered around the 40S-1 and 60S-1 sites respectively.^**26**,**27**^ Specifically, any ribosomal nucleotide or residue with at least one atom that was within 15 Å of any atom of C1237 of the 40S-1 site and the A2009 of the 60S-1 site respective was included in the fragment in its entirety.

The docking of ligands within the receptors was performed using the program rDock^**28**^, which was selected due to its ability to dock nucleic acids. The rbcavity command within rDoc identified empty spaces within the receptors using the two sphere method, with the small sphere set at a radius of 1.5 Å and the large sphere set at a radius of 4 Å. Only a single cavity, equivalent to an eRF1 pocket, was identified in each ribosomal subunit, corresponding to the 60S-1 and 40S-1 pockets, respectively.

The ligands selected for vitrtual screening to the 60S-1 pocket were acquired from the Drug Subset of the Cambridge Structural Database (CSD)^**21**^, encompassing all approved drugs worldwide listed in DrugBank and totaling approximately 17,400 compounds. This set of ligands was docked to the 60S-1 site receptor models using the dock_solv method within rDoc, which takes into account desolvation, and is recommended for docking nucleic acids. A total of 100 poses were generated for each ligand. The docking runs were completed using the University of Pennsylvania High Performance Computing (HPC) cluster, which uses the IBM Load Sharing Facility (LSF) environment. The library was split into 70 sub-libraries of approximately 250 compounds each, run in parallel to expedite the runs. After the docking run was completed, the compounds were ranked by the score of their best-scoring pose, and the compounds reflecting the top 1% of scores which were commercially available were selected for *in vitro* validation. This process led to the identification of telmisartan as a ligand for the 60S-1 pocket, and, by structural analogy, of candesartan.

Separately, candesartan cilexetil and N-Ac-RY-OMe were docked into the 60S-1 pocket and N-Ac-TS-OMe was docked into the 40S-1 pocket using rDoc as above. Then all 100 poses of these three compounds and of telmisartan and candesartan were rescored using the AnnapuRNA program^**29**^, with the top-scoring poses after the rescore chosen as the hypothesized model for binding into the 60S-1 or 40S-1 pocket. The ribosomal contacts with the each of the five TRIDs falling within 3.2 Å of the top rescored docked poses were identified using ChimeraX (see Fig. 4).

### Materials for PURE-LITE Assays

#### tRNAs and ribosome complexes

Atto647-ε-lysine–tRNA^Lys^ was prepared by labeling [^3^H]Lys-tRNA^Lys^ from *E. coli* with Atto647-NHS as previously described^**17**,**18**^. 80S–Stop-POST5 complexes were prepared from *Artemia salina* 80S ribosomes and Cricket Paralysis Virus (CrPV) IRES mRNA as previously described^**17**,**19**^. 80S–Stop-IRES concentrations were determined by A_260_ measurement. Stop-POST5 concentrations were determined from A_260_ measurements and independently estimated from the stoichiometry of radiolabeled pentapeptide associated with the 80S ribosomes. Typically, 22 ± 5% of the 80S ribosomes contained the pentapeptide. For Atto(pep)-Stop-POST5 complexes prepared using Atto647-labeled Lys-tRNA in place of unlabeled Lys-tRNA, Atto647 absorbance (A_647_) measurements were additionally used to determine POST5 concentration and were in close agreement with the concentration estimated from radiolabeled pentapeptide stoichiometry, consistent with previous reports^**17**,**18**^.

#### Reagents for Assays

Ataluren sodium salt was obtained as described^**19**^. Sartans were purchased from Med Chem Express. Sartan and ataluren concentrations were determined spectroscopically using the following extinction coefficients: Telmisartan, ε_296_ = 4.0 × 10^4^ M^−1^ cm^−1^; Candesartan, ε_254_ = 1.29 × 10^4^ M^−1^ cm^−1^, Candesartan cilexetil, ε_268_ = 3.16 × 10^4^ M^−1^ cm^−1^; Ataluren, ε_241_ = 3.22 × 10^4^ M^−1^ cm^−1^. Peptide and peptide derivatives were either purchased from Genscript or synthesized as described below. The concentrations of peptides containing His, Phe, Trp, or Tyr residues were determined by UV–Vis absorbance using their characteristic aromatic absorption, with approximate extinction coefficients of His ~200 M^−1^cm^−1^ at 215 nm, Phe ~200 M^−1^cm^−1^ at 257 nm, Tyr ~1,500 M^−1^cm^−1^ at 275 nm, and Trp ~5,500 M^−1^cm^−1^ at 280 nm; the concentrations of the remaining peptides, which lacked these UV-active residues, were calculated from their measured weights and compositions.

### PURE-LITE Assays

#### Termination assay

Termination assays were performed either at 25 °C or 37 °C in Buffer 4 (pH 7.4; 40 mM Tris HCl, pH 7.5, 80 mM NH_4_Cl, 5 mM Mg(Ac)2, 100 mM KOAc, and 3 mM β-mercaptoethanol). using a TECAN SPARK multimode plate reader equipped with a monochromator and either 96-well Greiner® or 384-well Corning black, flat-bottom plates, as described previously^**17**,**19**^. Two solutions were prepared in Buffer 4: (i) 0.05 μM Atto(pep)-Stop-POST5 and 1 mM GTP and (ii) eRF1 or the indicated eRF1 variant (0–0.5 μM), excess eRF3 (0.8 μM), and 1 mM GTP. Reactions were initiated by mixing the two solutions, with all concentrations reported as final concentrations after mixing. Peptide release was monitored continuously by measuring the decrease in fluorescence anisotropy resulting from dissociation of the Atto647-labeled pentapeptide following eRF1 or eRF1 variant-catalyzed hydrolysis of ribosome-bound pentapeptidyl-tRNA. For assays containing ataluren, Sartans, or peptides, the corresponding compounds were included in the RFC-containing solution. Time-dependent changes in fluorescence anisotropy were analyzed using GraphPad Prism and fit to a single-exponential model with an asymptote to determine peptide dissociation rates for Stop-POST5 complexes in the presence or absence of peptides, sartans, or ataluren. The resulting rate data were subsequently fit to the Michaelis–Menten equation to determine *k*_cat_ and *K*_M_ values for the release factor complex (RFC) and its variants. Here we note that the previously observed S-shaped dependence of termination rate on RFC concentration for ribosomes programmed with the reference CrPV-mRNA sequence (Fig. 1), measured at 25 °C^**17–19**^, was reversibly converted to a rectangular hyperbolic dependence for the measurements conducted at 37 °C in the present study,

#### Readthrough Assay

The readthrough assay measuring octapeptide formation from Stop-POST5 was performed as reported previously^**16**,**30**^, with the modification that [^3^H]-labeled signal of aa-tRNAs was used instead of [^35^S]-aa-Met-tRNA. Briefly, the Stop-IRES–POST5 complex (0.05 μM; 20 μL; Solution A) was preincubated in Buffer 4 at 37 °C for 2–3 min. In parallel, Solution B containing [^3^H]-Trp-tRNA, cognate Leu-tRNA and Met-tRNA (each 0.2 μM), eEF1A (1 μM), GTP (1 mM), and eEF2 (1 μM) was prepared in the absence or presence of increasing concentrations of peptides or Sartans. [^3^H]-Trp-tRNA served as the near-cognate tRNA for the UGA stop codon at codon 6 of the IRES sequence^**19**^. Readthrough was initiated by mixing Solutions A and B; all concentrations indicated above represent final concentrations after mixing. Reactions were incubated for 30 min at 37 °C, and then quenched with 150 μL ice-cold 0.5 M MES buffer (pH 6.0) and immediately placed on ice. Carrier 70S *E. coli* ribosomes (100 pmol) were added to aid in sedimentation, and the reaction mixtures were layered onto 1.1 M sucrose prepared in Buffer 4 (1.5X the reaction volume) and subjected to ultracentrifugation at 540,000 × g for 120 min at 4 °C. The resulting ribosome pellets were washed and dissolved in Buffer 4, and then analyzed by A_260_ measurement and quantification of the [^3^H]-labeled octapeptide (POST8)^**16**^ to determine readthrough efficiency.

### Dual-luciferase readthrough assay using rabbit reticulocyte lysate^31^

Dual-luciferase reporter mRNAs containing the PTC context in the vector pSGDlucV3.0 (Addgene item #119760) were synthesized in vitro and capped using the NEB Vaccinia Capping System (M2080S). The reporter constructs tested in this study contained four disease-associated premature termination codons (PTCs): G542X, R1162X, and W1282X in CFTR, which are associated with cystic fibrosis, and R2694X in FBN1, which is associated with Marfan syndrome. For each PTC reporter, a corresponding sense-control construct was generated in which the UGA PTC was replaced by the cognate UGG codon for tRNA^Trp^ while maintaining the identical surrounding patient sequence context of 75 nucleotides upstream and downstream of the PTC (Table S5). Synthetic DNA fragments were assembled using In-Fusion cloning (GenScript), and all constructs were sequence-verified across at least 200 nt of the relevant flanking regions before use. Capped reporter mRNAs were translated in nuclease-treated rabbit reticulocyte lysate (Promega, L4960) in the presence of vehicle or varying concentrations of test TRIDs, including ataluren, telmisartan, candesartan or candesartan cilexetil. Following translation reaction for 90 min, Firefly (Fluc) and Renilla (Rluc) luciferase activities were quantified on a TECAN Spark reader using the Promega Dual-Glo Luciferase Reporter Assay System (E1910) according to the manufacturer’s instructions (Promega). Readthrough efficiency was calculated from the Fluc/Rluc ratio and normalized to the corresponding matched sense control. Each condition was analyzed in technical triplicate with at least three biological replicates.

### Synthesis and purification of peptides and peptide derivatives

#### Resin Preparation

Peptide synthesis was performed on a 1.54–3.21 mmol scale using 2-chlorotrityl chloride resin (loading density: 1.00 mmol g^−1^). Prior to functionalization, the resin was transferred to a 60 mL solid-phase peptide synthesis (SPPS) vessel and swollen in anhydrous dichloromethane (DCM) for 1–2 min before solvent removal.

#### Loading of Fmoc-Protected Amino Acids

Fmoc-protected amino acids (1.0 equiv) were dissolved in anhydrous DCM containing DIPEA (3.0 equiv) and added to the pre-swollen resin. The mixture was stirred at room temperature for 30 min, followed by addition of MeOH (15 mL) and continued stirring for an additional 15 min. The reaction mixture was drained, and the resin was washed thoroughly with DMF. Subsequent removal of the Fmoc protecting group was carried out using the standard deprotection protocol described below.

#### Fmoc Deprotection

The resin was treated with 10 mL of 20% piperidine in DMF for 30 min at room temperature. The solution was drained and the deprotection step repeated once more. The resin was then washed with DMF (5X) prior to amino acid coupling.

#### Standard Amino Acid Couplings

Coupling reactions were performed by adding activated amino acid solutions to the deprotected resin. Each coupling mixture contained 1.5 equiv of the appropriate Fmoc-amino acid, 1.5 equiv HBTU, and 3.0 equiv DIPEA dissolved in DMF (10 mL). Solutions were pre-activated by swirling for 5–10 min before addition. The resin was stirred with the coupling mixture for 12 h at room temperature, then drained and washed.

#### N-terminal Acetylation

Following removal of the terminal Fmoc group, the resin was washed with DMF and treated with 6 mL of an acetylation solution consisting of Ac_2_O/DIPEA/DMF (1:2:3, v/v). The mixture was stirred for 30 min at room temperature.

#### Cleavage of the protected peptide

Upon completing all appropriate couplings, the resin was washed with 2x DMF and 3x DCM and then air-dried. To obtain the peptide with a free C-terminal acid, the resin was treated with a mild acidic cocktail (TFA/TIPS/DCM, 3:2:95, v/v/v). To obtain the peptide as a C-terminal methyl ester, the resin was instead exposed to MeOH/4 M HCl in dioxane (1:1). In both cases, the mixture was stirred for 30 min and concentrated under reduced pressure before purification.

#### Cyclization

Following cleavage of the peptide bearing a free N-terminal amine and a C-terminal methyl ester, the crude material was dissolved in DCM and treated with saturated aqueous NaHCO_3_. The biphasic mixture was stirred at room temperature overnight to promote intramolecular cyclization. The reaction mixture was then extracted with DCM (3x), and the combined organic layers were concentrated under reduced pressure to afford the cyclized product, RY-Cyclic and TS-Cyclic.

#### Purification

Crude peptides were purified by preparative reversed-phase HPLC using water/acetonitrile mobile phases containing 0.1% TFA. Elution was monitored at 214 nm, and collected fractions were analyzed by LC-MS. Fractions corresponding to the desired product were combined and lyophilized repeatedly to yield peptides as white solids. Final identity and purity were confirmed by MS, HPLC, and NMR spectroscopy.

### Generation, expression, and purification of eRF1 mutants

Human wild-type eRF1 was cloned into the pET-15b expression vector, and mutant constructs were generated by site-directed mutagenesis using the primers listed in Table S2. All mutations were confirmed by whole-plasmid sequencing using the Plasmidsaurus Oxford Nanopore sequencing platform (R10.4.1). Wild-type and mutant eRF1 proteins were expressed in *Escherichia coli* BL21(DE3) CodonPlus cells following induction with 0.5 mM IPTG at 16 °C for 12–16 h. Proteins were purified by TALON cobalt-affinity chromatography (Takara Bio, USA) and eluted with up to 500 mM imidazole. Purified proteins were desalted into storage buffer containing 50 mM HEPES, 100 mM KCl, 10% glycerol, and 1 mM DTT (pH 8.0), aliquoted, flash-frozen in liquid nitrogen, and stored at −80 °C. Protein expression and purification were performed as previously described^**32**^, with minor modifications, including expression at 16 °C rather than 25 °C, use of 100 mM rather than 50 mM KCl, omission of EDTA, and elution with up to 500 mM rather than 250 mM imidazole.

#### Variant characterization

The sequences of 32 eRF1 variants potentially important for binding to the ribosome were verified by Plasmidsaurus Oxford Nanopore sequencing platform (R10.4.1). Wild-type and mutant eRF1 proteins were expressed in *Escherichia coli* BL21(DE3) CodonPlus cells and purified by TALON Co^2^+-affinity chromatography (Takara Bio, USA). SDS–PAGE analysis (Fig. S1) showed that wild-type and variant eRF1 purified preparations exclusively or predominantly contained a single protein band.. corresponding to a mass of approximately 51 kDa as determined by MALDI-TOF mass spectrometry and expected for intact eRF1 with an N-terminal containing oligohisitidine.

#### MALDI-TOF mass spectrometry

Purified wild-type and mutant eRF1 proteins were diluted to 15 µM in 10 mM HEPES (pH 7.5). The sinapinic acid matrix was prepared at 10 mg/mL in H_2_O, and the protein and matrix solutions were mixed at a 1:1 volume ratio^**33**^. The mixture was spotted onto a MALDI target plate and allowed to air-dry. Intact-protein mass spectra were acquired using a rapifleX MALDI-TOF/TOF mass spectrometer (Bruker Daltonics) operated in positive-ion linear mode.

## Acknowledgements

This work was supported by grants to BSC (NIH R01-GM-127374, Cystic Fibrosis Foundation COOPER26GO, Penn Institute for RNA Innovation) and DMC (Internal and discretionary funds at the University of Pennsylvania).

## References

1. Boycott KM, Lau LP, Cutillo CM, Austin CP. International collaborative actions and transparency to understand, diagnose, and develop therapies for rare diseases. EMBO Molecular Medicine. 2019 May 1;11(5):EMMM201910486.

2. M. Mort, D. Ivanov, D. N. Cooper, N. A. Chuzhanova, A meta-analysis of nonsense mutations causing human genetic disease. Hum. Mutat. 29, 1037–1047 (2008)

3. Martins-Dias P, Romão L. Nonsense suppression therapies in human genetic diseases. Cellular and Molecular Life Sciences. 2021 May;78(10):4677–701.

4. Morais, P., Zhang, R., & Yu, Y. T. (2024). Therapeutic Nonsense Suppression Modalities: From Small Molecules to Nucleic Acid-Based Approaches. Biomedicines, 12(6), 1284

5. Torices L, Nunes-Xavier CE, Pulido R. Therapeutic Potential of Translational Readthrough at Disease-Associated Premature Termination Codons From Tumor Suppressor Genes. IUBMB life. 2025 May;77(5):e70018.

6. Sharma, J., Du, M., Wong, E., Mutyam, V., Li, Y., Chen, J., … & Bedwell, D. M. (2021). A small molecule that induces translational readthrough of CFTR nonsense mutations by eRF1 depletion. Nature communications, 12(1), 4358.

7. Gurzeler LA, Link M, Ibig Y, Schmidt I, Galuba O, Schoenbett J, Gasser-Didierlaurant C, Parker CN, Mao X, Bitsch F, Schirle M. Drug-induced eRF1 degradation promotes readthrough and reveals a new branch of ribosome quality control. Cell reports. 2023 Sep 26;42(9).

8. Baradaran-Heravi, A., Balgi, A. D., Hosseini-Farahabadi, S., Choi, K., Has, C., & Roberge, M. (2021). Effect of small molecule eRF3 degraders on premature termination codon readthrough. Nucleic acids research, 49(7), 3692–3708.

9. Ko W, Porter JJ, Spelier S, Sorensen EG, Bhatt P, Gabell JT, van der Windt I, Couch T, Coote K, Mense M, Beekman JM. ACE-tRNAs are a platform technology for suppressing nonsense mutations that cause cystic fibrosis. Nucleic acids research. 2025 Jul 22;53(13):gkaf675.

10. Pierce SE, Erwood S, Oye K, An M, Krasnow N, Zhang E, Raguram A, Seelig D, Osborn MJ, Liu DR. Prime editing-installed suppressor tRNAs for disease-agnostic genome editing. Nature. 2025 Dec 4;648(8092):191–202.ACE-tRNA

11. Chen J, Zhou M, Dong S, Gong F, Tennakoon R, Seto BY, Chen ZR, Zhou ZP, Pan J, Xu Y, Luozhong S. Nonviral delivery of chemically modified tRNA rescues nonsense mutations in cystic fibrosis. Science. 2026 Aug 27;393(6814):eaeb0054.

12. Zhu L, Muhtar P, Goyama S, Yoshimi A. Advances in antisense oligonucleotide treatment for cancer. Japanese Journal of Clinical Oncology. 2026 Feb 3:hyag017.

13. Michorowska S. Ataluren—Promising Therapeutic Premature Termination Codon Readthrough Frontrunner. Pharmaceuticals. 2021 Aug 9;14(8):785.

14. Manjunath R, Gaonkar SL. Ataluren, a 1, 2, 4-Oxadiazole-Containing Drug Candidate for Nonsense Mutations: Major Breakthroughs and an Up-to-Date Process Chemistry Review. ChemistrySelect. 2025;10(2).

15. Nagel-Wolfrum K, Wenck N, Zorin M. Translational readthrough using TRIDs-Achievements and challenges for the treatment of inherited retinal disorders. Molecular Aspects of Medicine. 2026 Apr 1;108:101455.

16. Ng, M. Y., Li, H., Ghelfi, M. D., Goldman, Y. E., & Cooperman, B. S. (2021). Ataluren and aminoglycosides stimulate read-through of nonsense codons by orthogonal mechanisms. Proceedings of the National Academy of Sciences, 118(2), e2020599118.

17. Huang, S., Bhattacharya, A., Ghelfi, M. D., Li, H., Fritsch, C., Chenoweth, D. M., … & Cooperman, B. S. (2022). Ataluren binds to multiple protein synthesis apparatus sites and competitively inhibits release factor-dependent termination. Nature Communications, 13(1), 2413.

18. Ghelfi MD, Bhat SY, Li H, Cooperman BS. A high-throughput assay for in vitro determination of release factor-dependent peptide release from a pretermination complex by fluorescence anisotropy—application to nonsense suppressor screening and mechanistic studies. Biomolecules. 2023 Jan 27;13(2):242.

19. Bhat, S. Y., Bhattacharya, A., Li, H., Cui, X., Lueck, J. D., Goldman, Y. E., & Cooperman, B. S. (2025). Mechanism-based approach in designing patient-specific combination therapies for nonsense mutation diseases. Nucleic Acids Research, 53(6), gkaf216.

20. Chen Y, Tian M, Zhu Z, Qian Y, Xu Y, Gong P, Church G, Liu C. Engineered translation release factor 1 suppresses disease-causing nonsense mutations and enables multi-site nonstandard amino acids incorporation. bioRxiv. 2025 Dec 25:2025-12.

21. Bryant, M. J.; Black, S. N.; Blade, H.; Docherty, R.; Maloney, A. G. P.; Taylor, S. C. The CSD Drug Subset: The Changing Chemistry and Crystallography of Small Molecule Pharmaceuticals. Journal of Pharmaceutical Sciences 2019, 108 (5), 1655–1662. https://doi.org/10

22. Bezzerri, V, Pegoraro, A, Hristodor, AM, Crane, GM, MeneghelliI Brignole. C et al. Ataluren improves hematopoietic and pancreatic disorders in Shwachman-Diamond syndrome patients: a compassionate program case-series. Nat Commun. (2025) 16:8189. doi: 10.1038/s41467-025-63137-321.

23. Lentini, L., Perriera, R., Corrao, F., Melfi, R., Tutone, M., Carollo, P. S., … & Pibiri, I. (2025). A precision medicine approach to primary immunodeficiency disease: Ataluren strikes nonsense mutations once again. Molecular Therapy, 33(7), 3231–3241.

24. Shao, S.; Murray, J.; Brown, A.; Taunton, J.; Ramakrishnan, V.; Hegde, R. S. Decoding Mammalian Ribosome-mRNA States by Translational GTPase Complexes. Cell 2016, 167 (5), 1229–1240.e15. 10.1016/j.cell.2016.10.046.

25. Powers, K. T.; Flint Stevenson-Jones Yadav, N.; Amthor, B.; Bufton, J. C.; Ufuk Borucu Shen, D.; Becker, J. P.; Lavysh, D.; Hentze, M. W.; Kulozik, A. E.; Neu-Yilik, G.; Schaffitzel, C. Blasticidin S Inhibits Mammalian Translation and Enhances Production of Protein Encoded by Nonsense mRNA. Nucleic Acids Research 2021, 49 (13), 7665–7679. 10.1093/nar/gkab532.

26. Bhaskar, V.; Graff-Meyer, A.; Schenk, A. D.; Cavadini, S.; von Loeffelholz, O.; Natchiar, S. K.; Artus-Revel, C. G.; Hotz, H.-R.; Bretones, G.; Klaholz, B. P.; Chao, J. A. Dynamics of uS19 C-Terminal Tail during the Translation Elongation Cycle in Human Ribosomes. Cell Reports 2020, 31 (1), 107473. 10.1016/j.celrep.2020.03.037.

27. Voorhees, R. M.; Fernández, I. S.; Scheres, S. H. W.; Hegde, R. S. Structure of the Mammalian Ribosome-Sec61 Complex to 3.4 Å Resolution. Cell 2014, 157 (7), 1632–1643. 10.1016/j.cell.2014.05.024.

28. Ruiz-Carmona, S.; Alvarez-Garcia, D.; Foloppe, N.; Garmendia-Doval, A. B.; Juhos, S.; Schmidtke, P.; Barril, X.; Hubbard, R. E.; Morley, S. D. rDock: A Fast, Versatile and Open Source Program for Docking Ligands to Proteins and Nucleic Acids. PLoS Computational Biology 2014, 10 (4), e1003571. 10.1371/journal.pcbi.1003571.

29. Stefaniak, F.; Bujnicki, J. M. AnnapuRNA: A Scoring Function for Predicting RNA-Small Molecule Binding Poses. PLOS Computational Biology 2021, 17 (2), e1008309. 10.1371/journal.pcbi.1008309

30. Ng MY, Zhang H, Weil A, Singh V, Jamiolkowski R, Baradaran-Heravi A, Roberge M, Jacobson A, Friesen W, Welch E, Goldman YE. New in vitro assay measuring direct interaction of nonsense suppressors with the eukaryotic protein synthesis machinery. ACS Medicinal Chemistry Letters. 2018 Nov 21;9(12):1285.

31. Loughran, G., Li, X., O’Loughlin, S., Atkins, J.F. and Baranov, P.V., 2023. Monitoring translation in all reading frames downstream of weak stop codons provides mechanistic insights into the impact of nucleotide and cellular contexts. Nucleic acids research, 51(1), pp.304–314.

32. Ito, L. Frolova, A. Seit-Nebi, A. Karamyshev, L. Kisselev, & Y. Nakamura, Omnipotent decoding potential resides in eukaryotic translation termination factor eRF1 of variant-code organisms and is modulated by the interactions of amino acid sequences within domain 1, Proc. Natl. Acad. Sci. U.S.A. 99 (13) 8494–8499, 10.1073/pnas.142690099 (2002).

33. Giménez E, Benavente F, Barbosa J, Sanz-Nebot V. Ionic liquid matrices for MALDI-TOF-MS analysis of intact glycoproteins. Analytical and bioanalytical chemistry. 2010 Sep;398(1):357–65

