## Supplemental Tables and Figures for "Mechanism-Based Discovery of New TRIDs Targeting Ribosome Pockets"

### **Supplementary Information**

#### **Contents**

- **1. Tables S1 – S6**
- **2. Figures S1 and S2**

**Table S1.** Steady-state kinetic parameters ($K_m$, $k_{cat}$, and $k_{cat}/K_m$) for peptide release catalyzed by wild-type and mutant human eRF1 proteins. Values are presented as mean ± SD.

| Serial # | eRF1 mutant | $K_m$, nM | $k_{cat}$, min<sup>-1</sup> | $k_{cat}/K_m$, $\mu\text{M}^{-1}\text{min}^{-1}$ |
| --- | --- | --- | --- | --- |
| 1 | WT | 10 ± 2 | 0.30 ± 0.02 | 30 ± 6 |
| 2 | T76A | 104 ± 11 | 0.21 ± 0.01 | 2.1 ± 0.2 |
| 3 | S77A | 13 ± 1 | 0.71 ± 0.02 | 54.6 ± 4.5 |
| 4 | S77E | 51 ± 8 | 0.19 ± 0.02 | 3.7 ± 0.7 |
| 5 | Q80A | 77 ± 10 | 0.18 ± 0.01 | 2.3 ± 0.3 |
| 6 | G98N | 32.8 ± 6.2 | 0.26 ± 0.01 | 8.1 ± 1.6 |
| 7 | R414A | 85 ± 24 | 0.22 ± 0.04 | 2.6 ± 0.8 |
| 8 | Y415A | 12 ± 1 | 0.17 ± 0.01 | 14.2 ± 1.4 |
| 9 | Y415W | 9.5 ± 4.1 | 0.18 ± 0.01 | 20.2 ± 8.8 |
| 10 | Q185A | 9.8 ± 0.6 | 0.08 ± 0.01 | 0.82 ± 0.06 |
| 11 | M314A | 5.2 ± 2.8 | 0.15 ± 0.02 | 28.4 ± 12.2 |
| 12 | M314L | 2.8 ± 0.6 | 0.13 ± 0.01 | 46.4 ± 10.5 |
| 13 | G216N | 4.3 ± 0.7 | 0.15 ± 0.01 | 34.7 ± 6.7 |
| 14 | Y424A | 9.6 ± 1.7 | 0.17 ± 0.01 | 17.6 ± 3.2 |
| 15 | G105V | 9.5 ± 2.7 | 0.18 ± 0.01 | 18.9 ± 5.5 |
| 16 | G105N | 6.1 ± 1.9 | 0.18 ± 0.01 | 30.6 ± 10.4 |
| 17 | V417N | 19.4 ± 4.7 | 0.14 ± 0.01 | 7.2 ± 1.8 |
| 18 | S215E | 4.2 ± 0.9 | 0.13 ± 0.01 | 32.5 ± 7.2 |
| 19 | G386N | 7.8 ± 1.7 | 0.13 ± 0.01 | 16.5 ± 4.1 |
| 20 | E418A | 10.5 ± 2.3 | 0.20 ± 0.01 | 18.6 ± 4.2 |
| 21 | E423A | 6.9 ± 1.2 | 0.16 ± 0.01 | 22.5 ± 4.1 |
| 22 | L193N | 33.7 ± 10.1 | 0.10 ± 0.01 | 2.9 ± 0.9 |
| 23 | T388A | 57.8 ± 10.8 | 0.11 ± 0.02 | 2.1 ± 0.8 |
| 24 | T388E | 8.5 ± 1.4 | 0.31 ± 0.02 | 37.3 ± 6.6 |
| 25 | A387N | 23.2 ± 1.7 | 0.14 ± 0.01 | 6.1 ± 0.6 |
| 26 | E104A | 20 ± 3 | 0.27 ± 0.03 | 13.5 ± 2.5 |
| 27 | E313A | 22.9 ± 5.4 | 0.26 ± 0.04 | 11.2 ± 3.2 |
| 28 | H199A | 34.9 ± 6.8 | 0.12 ± 0.02 | 3.6 ± 0.8 |
| 29 | N200A | 31 ± 5 | 0.12 ± 0.01 | 3.9 ± 0.7 |
| 30 | R416A | 41.4 ± 9.1 | 0.24 ± 0.02 | 5.9 ± 1.4 |
| 31 | R189A | N.D | ~0.02 | N.D |
| 32 | R192A | N.D | ~0.02 | N.D |

**Table S2: List of Forward and Reverse Primers Used for Site-Directed Mutagenesis of Human eRF1**

| Mutants | Forward Primers (5'-3') | Reverse Primers (5'-3') |
| --- | --- | --- |
| T76A | GTTGTACAGATGTAATGGCTCCC | GTTGTACAGATGTAATGGCTCCC |
| S77A | GAGTCTTTGTTGTACAGATGTAATGGCT | GAGTCTTTGTTGTACAGATGTAATGGCT |
| S77E | AGCCATTACAGAAGTACAACAAAGACTC | CCCAGGACTGAAAGG |
| Q80A | ATCTGTACAAGCGAGACTCAAACTTTATAACAAAG | CTTTGTTATAAAGTTTGAGTCTCGCTTGTACAGAT |
| G98N | TGTATACTGTAACACAATTGTAACAGAAG | ACCAGACCATTTGGAG |
| E104A | TGTAACAGAAGCGGGAAAGGAAAAGAAAG | ATTGTTCCACAGTATACAAC |
| G105N | AACAGAAGAAAACAAGGAAAAGAAAGTCAAC | ACAATTGTTCCACAGTATAC |
| G105V | AACAGAAGAAGTGAAGGAAAAGAAAGTC | ACAATTGTTCCACAGTATAC |
| Q185A | TAGAGGAGGTGCGTCAGCCTTGC | GCAAGGCTGACGCACCTCCTCTA |
| R189A | GTCAGCCTTGGCGTTTGCCCGTTTAAG | CTTAAACGGGCAAACGCCAAGGCTGAC |
| R192A | GCGTTTTGCCGCGTTAAGAATGGAAAAG | CTTTTCCATTCTTAACGCGGCAAAACGC |
| L193N | TTTTGCCCGTAACAGAATGGAAAAGC | CGCAAGGCTGACTGAC |
| H199A | GGAAAAGCGAGCGAACTATGTTCGGAAAG | ATTCTTAAACGGGCAAAAC |
| N200A | AAAGCGACATGCGTATGTTCGGAAAGTAG | TCCATTCTTAAACGGG |
| S215E | GCTGTTTATTGAAGGGGACAAAGTG | TGCACAGCAGTCTCTG |
| G216N | GTTTATTTCTAACGACAAAGTGAATGTGGC | AGCTGCACAGCAGTC |
| E313A | AAAGGCTTTGGCGATGGGAGCTG | AGTGTATCTTCAACGCC |
| M314A | GGCTTTGGAAGCGGGAGCTGTAG | TTTAGTGTATCTTCAACGC |
| M314L | GGCTTTGGAACTGGGAGCTGTAG | TTTAGTGTATCTTCAACGCCAAAAC |
| G386N | TAAAAAATTTAACGCTACGTTGGAAATTGTC | TAGTTGTTAGCAAACCATTC |
| A387N | AAAATTTGGAAACACGTTGGAAATTGTCAC | TTATAGTTGTTAGCAAACC |
| T388A | ATTTGGAGCTGCGTTGGAAATTG | TTTTTATAGTTGTTAGCAAACC |
| T388E | ATTTGGAGCTGAATTGGAAATTGTCACAG | TTTTTATAGTTGTTAGCAAACC |
| T388N | ATTTGGAGCTAACTTGGAAATTGTCACAGATAAATC | TTTTTATAGTTGTTAGCAAACC |
| R414A | AGGTATCTTGGCGTACCGAGTAG | CCAATTCCACCAAATCC |
| Y415A | TATCTTGCGGGCGCGAGTAGATTTCCAG | CCTCCAATTCCACCAAATC |
| Y415W | TATCTTGCGGTGGCGAGTAGATTTC | GAAATCTACTCGCCACCGCAAGATA |
| R416A | CTTGCGGTACGCGGTAGATTTCCAG | ATACCTCCAATTCCACC |
| V417N | GCGGTACCGAAACGATTTCCAGG | AAGATACCTCCAATTCCAC |
| E418A | GTACCGAGTAGCGTTCCAGGGAATG | CGCAAGATACCTCCAATTC |
| E423A | CCAGGGAATGGCGTACCAAGGAG | AAATCTACTCGGTACCG |
| Y424A | GGGAATGGAAGCGCAAGGAGGAGACG | TGGAAATCTACTCGGTAC |

| Table S3. 28S rRNA and rPL40 contacts with TRIDs within 60S-1 |  |  |  |  |
| --- | --- | --- | --- | --- |
| 60S-1 contact | N-Ac-RY-OMe | Telmisartan | Candesartan | Candesartan Cilexetil |
| G 1981 | +++ | ++ | + | ++ |
| G 1982 | * | + | +++ | + |
| A 1983 | + | ++ | ++ | +++ |
| U 1986 | * | ++ |  |  |
| C 1987 | + | +++ | +++ |  |
| G 1988 | * |  |  |  |
| U 2006 | +++ | * |  | + |
| G 2007 | +++ | ++ | + | ++ |
| U 2008 | ++ | +++ | ++ | +++ |
| A 2009 | ++ | +++ | * | +++ |
| A 2010 | * | ++ | * | +++ |
| U 4419 | ++ | + | ++ | +++ |
| U 4420 | ++ | ++ | +++ | ++ |
| LYS 128 |  |  | * | ++ |
| Distance coding, in Å: ≤2.0 , +++; >2.0–2.3, ++; >2.3–2.6 +; >2.6–3.2, * |  |  |  |  |

| Table S4. 18S rRNA Contacts with N-Ac-TS-OMe within 40S-1 |  |
| --- | --- |
| 40S-1 contact | distance<sup>a</sup> |
| C 1234 | ++ |
| G 1235 | ++ |
| G 1236 | + |
| C 1237 | ++ |
| U 1238 | * |
| U 1242 | ++ |
| U 1243 | * |
| U 1244 | * |
| C 1518 | +++ |
| U 1519 | + |
| G 1520 | ++ |
| C 1521 | * |
| <sup>a</sup>Distance coding, in Å: ≤2.0 , +++; >2.0–2.3, ++; >2.3–2.6 +; >2.6–3.2, * |  |

**Table S5.PTC Sequence contexts used for rabbit reticulocyte extract experiments**

G542X TGA:

TAC AGA AGC GTC ATC AAA GCA TGC CAA CTA GAA GAG GAC ATC TCC AAG
TTT GCA GAG AAA GAC AAT ATA GTT CTT TGA GAA GGT GGA ATC ACA CTG
AGT GGA GGT CAA CGA GCA AGA ATT TCT TTA GCA AGA GCA GTA TAC AAA
GAT GCT GAT

W1282X TGA:

TTT TTG AGA CTA CTG AAC ACT GAA GGA GAA ATC CAG ATC GAT GGT GTG
TCT TGG GAT TCA ATA ACT TTG CAA CAG TGA AGG AAA GCC TTT GGA GTG
ATA CCA CAG AAA GTA TTT ATT TTT TCT GGA ACA TTT AGA AAA AAC TTG
GAT CCC TAT

R1162X TGA:

ATG AAT ATC ATG AGT ACA TTG CAG TGG GCT GTA AAC TCC AGC ATA GAT
GTG GAT AGC TTG ATG CGA TCT GTG AGC TGA GTC TTT AAG TTC ATT GAC
ATG CCA ACA GAA GGT AAA CCT ACC AAG TCA ACC AAA CCA TAC AAG AAT
GGC CAA CTC

R2694X TGA:

GGT TAC CTG TGT GGC TGT CCA CCT GGT TAC TTC CGC ATA GGC CAA GGG
CAC TGT GTT TCT GGA ATG GGC ATG GGC TGA GGA AAC CCA GAG CCA CCT
GTC AGT GGT GAA ATG GAT GAC AAT TCA CTC TCC CCA GAG GCT TGT TAC
GAG TGT AAG

| **Table S6. Examples of Positive Ataluren Effects on Readthrough of PTC Diseases Reported on Humans, Animals , and/or Cells Isolated from Patients** |  |  |
| --- | --- | --- |
| Mutated protein | Disease | References |
| ABCC6 | Pseudoxanthoma Elasticum | 1 |
| BMPR2 | HPAH | 2,3 |
| CFTR | CF | 4-8 |
| Dysferlin | Dysferlinopathy | 9,10 |
| Dystrophin | DMD | 11 |
| FANCC | Fanconi anemia | 12 |
| FAT1 | Cancer | 13 |
| HERG | Long QT syndrome | 14 |
| LRBA | Lipopolysaccharide responsive Beige anchor | 15 |
| MCIDAS | primary ciliary dyskinesia (PCD) | 16 |
| MERTK | retinitis pigmentosa | 17 |
| NBCe1 | renal tubular acidosis | 18 |
| NF1 | Several diseases | 19 |
| Notch1 | HNSCC | 13 |
| PAX6 | Congenital aniridia | 20,21 |
| RP2 | xeroderma pigmentosum | 22 |
| SBDS | Shwachman-Diamond Syndrome | 23 |
| SMAD9 | HPAH | 3 |
| USH1C | USHER Syndrome | 24 |
| USH2A | USHER Syndrome | 25 |
| WRN | Werner syndrome | 26 |

#### References for Table S6

1. Zhou, Y., Jiang, Q., Takahagi, S., Shao, C., & Uitto, J. (2013). Premature termination codon read-through in the ABCC6 gene: potential treatment for pseudoxanthoma elasticum. *Journal of Investigative Dermatology*, 133(12), 2672-2677.
2. Long, L., Yang, X., Southwood, M., Moore, S., Crosby, A., Upton, P. D., ... & Morrell, N. W. (2020). Targeting translational read-through of premature termination mutations in BMPR2 with PTC124 for pulmonary arterial hypertension. *Pulmonary Circulation*, 10(3), 2045894020935783.
3. Drake, K. M., Dunmore, B. J., McNelly, L. N., Morrell, N. W., & Aldred, M. A. (2013). Correction of nonsense BMPR2 and SMAD9 mutations by ataluren in pulmonary arterial hypertension. *American journal of respiratory cell and molecular biology*, 49(3), 403-409.
4. Welch, E. M., Barton, E. R., Zhuo, J., Tomizawa, Y., Friesen, W. J., Trifillis, P., ... & Sweeney, H. L. (2007). PTC124 targets genetic disorders caused by nonsense mutations. *Nature*, 447(7140), 87-9

5. Sermet-Gaudelus, I., Boeck, K. D., Casimir, G. J., Vermeulen, F., Leal, T., Mogenet, A., ... & Miller, L. L. (2010). Ataluren (PTC124) induces cystic fibrosis transmembrane conductance regulator protein expression and activity in children with nonsense mutation cystic fibrosis. *American journal of respiratory and critical care medicine*, 182(10), 1262-1272.
6. Du, M., Liu, X., Welch, E. M., Hirawat, S., Peltz, S. W., & Bedwell, D. M. (2008). PTC124 is an orally bioavailable compound that promotes suppression of the human CFTR-G542X nonsense allele in a CF mouse model. *Proceedings of the National Academy of Sciences*, 105(6), 2064-2069.
7. Pibiri, I., Lentini, L., Melfi, R., Gallucci, G., Pace, A., Spinello, A., ... & Di Leonardo, A. (2015). Enhancement of premature stop codon readthrough in the CFTR gene by Ataluren (PTC124) derivatives. *European journal of medicinal chemistry*, 101, 236-24
8. Lentini, L., Melfi, R., Cancemi, P., Pibiri, I., & Di Leonardo, A. (2019). Caffeine boosts Ataluren's readthrough activity. *Heliyon*, 5(6).
9. Wang, B., Yang, Z., Brisson, B. K., Feng, H., Zhang, Z., Welch, E. M., ... & Sweeney, H. L. (2010). Membrane blebbing as an assessment of functional rescue of dysferlin-deficient human myotubes via nonsense suppression. *Journal of applied physiology*, 109(3), 901-905.
10. Seo, K., Kim, E. K., Choi, J., Kim, D. S., & Shin, J. H. (2021). Functional recovery of a novel knockin mouse model of dysferlinopathy by readthrough of nonsense mutation. *Molecular Therapy Methods & Clinical Development*, 21, 702-709.
11. Finkel, R. S., Flanigan, K. M., Wong, B., Bönnemann, C., Sampson, J., Sweeney, H. L., ... & Peltz, S. W. (2013). Phase 2a study of ataluren-mediated dystrophin production in patients with nonsense mutation Duchenne muscular dystrophy. *PloS one*, 8(12), e81302.
12. Hristodor, A. M., Cappelli, E., Baldisseri, E., Valli, R., Montalbano, G., Micheloni, G., ... & Bezzerri, V. (2025). Development of translational read-through-inducing drugs as novel therapeutic options for patients with Fanconi anemia. *Cell Death Discovery*, 11(1), 286.
13. Wu, M. H., Lu, R. Y., Yu, S. J., Tsai, Y. Z., Lin, Y. C., Bai, Z. Y., ... & Cai, B. H. (2022). PTC124 rescues nonsense mutation of two tumor suppressor genes NOTCH1 and FAT1 to repress HNSCC cell proliferation. *Biomedicine*, 10(11), 2948.
14. Yu, H., Liu, X., Huang, J., Zhang, Y., Hu, R., & Pu, J. (2014). Comparison of read-through effects of aminoglycosides and PTC124 on rescuing nonsense mutations of HERG gene associated with long QT syndrome. *International journal of molecular medicine*, 33(3), 729-735
15. Lentini, L., Perriera, R., Corrao, F., Melfi, R., Tutone, M., Carollo, P. S., ... & Pibiri, I. (2025). A precision medicine approach to primary immunodeficiency disease: Ataluren strikes nonsense mutations once again. *Molecular Therapy* 2025 Jul 2;33(7):3231-41.
16. Lee, D. D. H., Cardinale, D., Nigro, E., Butler, C. R., Rutman, A., Fassad, M. R., ... & O'Callaghan, C. (2021). Higher throughput drug screening for rare respiratory diseases: readthrough therapy in primary ciliary dyskinesia. *European Respiratory Journal*, 58(4).

17. Ramsden, C. M., Nommiste, B., R. Lane, A., Carr, A. J. F., Powner, M. B., JK Smart, M., ... & Coffey, P. J. (2017). Rescue of the MERTK phagocytic defect in a human iPSC disease model using translational read-through inducing drugs. *Scientific reports*, 7(1), 51.
18. Fang, Y. W., Yang, S. S., Chau, T., Nakamura, M., Yamazaki, O., Seki, G., ... & Lin, S. H. (2015). Therapeutic effect of prenatal alkalization and PTC124 in Na<sup>+</sup>/HCO<sub>3</sub><sup>-</sup>-cotransporter 1 p. W516\* knock-in mice. *Gene therapy*, 22(5), 374-381.
19. Wu, C., Iyer, S., Wolfe, S. A., & Jacobson, A. (2022). Functional restoration of mouse Nf1 nonsense alleles in differentiated cultured neurons. *Journal of Human Genetics*, 67(11), 661-668.
20. Cunha, D. L., Sarkar, H., Eintracht, J., Harding, P., Zhou, J. H., & Moosajee, M. (2023). Restoration of functional PAX6 in aniridia patient iPSC-derived ocular tissue models using repurposed nonsense suppression drugs. *Molecular Therapy-Nucleic Acids*, 33, 240-253.
21. Liu, X., Zhang, Y., Zhang, B., Gao, H., & Qiu, C. (2020). Nonsense suppression induced readthrough of a novel PAX6 mutation in patient-derived cells of congenital aniridia. *Molecular Genetics & Genomic Medicine*, 8(5), e1198
22. Schwarz, N., Carr, A. J., Lane, A., Moeller, F., Chen, L. L., Aguilà, M., ... & Hardcastle, A. J. (2015). Translational read-through of the RP2 Arg120stop mutation in patient iPSC-derived retinal pigment epithelium cells. *Human molecular genetics*, 24(4), 972-986.
23. Bezzerri, V., Lentini, L., Api, M., Busilacchi, E. M., Cavalieri, V., Pomilio, A., ... & Cipolli, M. (2022). Novel Translational Read-through-Inducing Drugs as a Therapeutic Option for Shwachman-Diamond Syndrome. *Biomedicines*, 10(4), 886
24. T. Goldmann et al., A Comparative Evaluation of NB30, NB54 And PTC124 In Translational Read-Through Efficacy for Treatment of an USH1C Nonsense Mutation. *EMBO Mol. Med.* **4**, 1186-1199 (2012).
25. Samanta, A., Stingl, K., Kohl, S., Ries, J., Linnert, J., & Nagel-Wolfrum, K. (2019). Ataluren for the treatment of usher syndrome 2A caused by nonsense mutations. *International journal of molecular sciences*, 20(24), 6274.
26. Agrelo, R., Sutz, M. A., Setien, F., Aldunate, F., Esteller, M., Da Costa, V., & Achenbach, R. (2015). A novel Werner Syndrome mutation: pharmacological treatment by read-through of nonsense mutations and epigenetic therapies. *Epigenetics*, 10(4), 329-341.

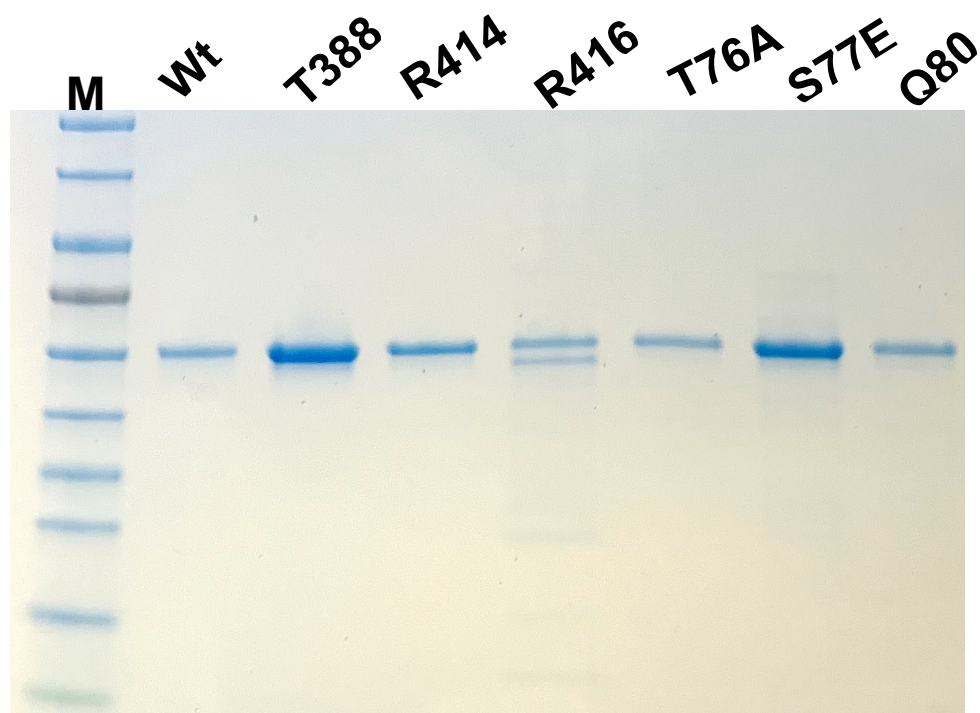

**Fig. S1** SDS-PAGE of eRF1 variants presented in Table 2

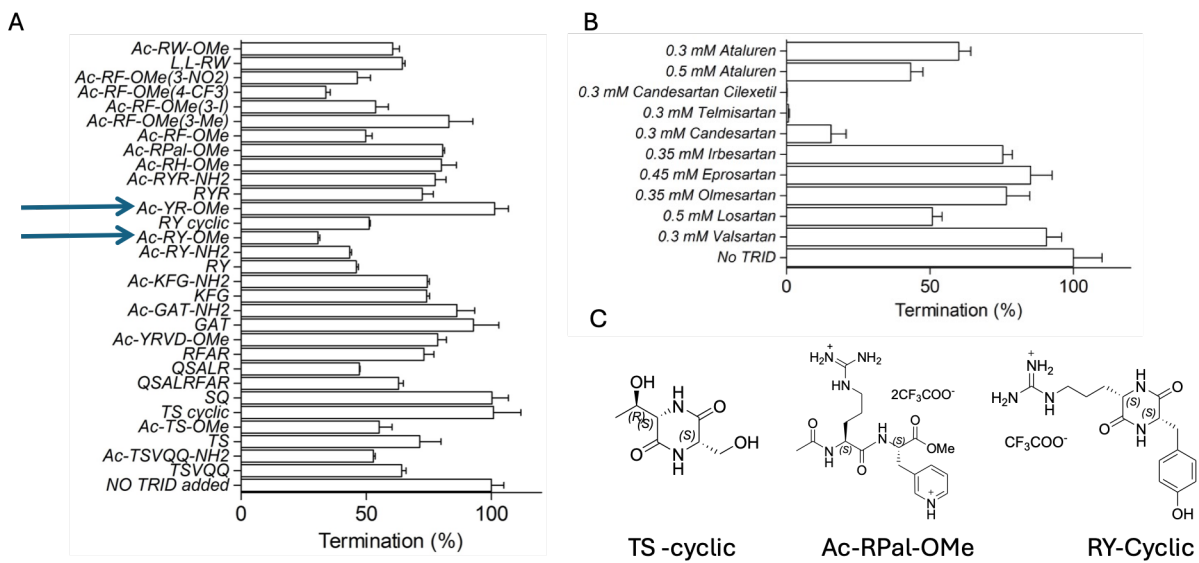

**Figure S2. Termination activities of peptide analogs and sartans.**

**A. Peptides.** Termination activities of the indicated peptides, represented using one-letter amino acid codes; phenylalanine (F) substitutions are indicated in parentheses. Peptides were tested at an approximate final concentration of 0.75 mM. Arrows highlight the higher activity of N-Ac-RY-OMe (RY-1), which mimics the R414–Y415 sequence within eRF1, compared with Ac-YR-OMe peptide. Pal in Ac-RPal-OMe denotes 3-(pyridyl)alanine.

**B. Sartans.** Termination activities of the indicated sartans. The concentration of each compound is indicated above the corresponding bar. Data are presented as mean ± SD from three determinations.

**C.** Structures of TS cyclic, Ac-RPal-OMe and RY-cyclic.
